# The 1/f-like behavior of neural field spectra are a natural consequence of noise driven brain dynamics

**DOI:** 10.1101/2023.03.10.532077

**Authors:** Mark A. Kramer, Catherine J. Chu

## Abstract

Consistent observations across recording modalities, experiments, and neural systems find neural field spectra with 1/f-like scaling, eliciting many alternative theories to explain this universal phenomenon. We show that a general dynamical system with stochastic drive and minimal assumptions generates 1/f-like spectra consistent with the range of values observed *in vivo*, without requiring a specific biological mechanism or collective critical behavior.

## INTRODUCTION

Transient oscillations are a prominent feature of macroscopic neural field activity [1], linked to brain function [2] and dysfunction [3], [4]. Oscillations appear as narrowband increases in the spectrum above an aperiodic background in which the power *P* decreases proportional to the frequency *f* raised to an exponent *β*: *P* ∝ *f*^*β*^. Characterizing oscillations while accounting for the aperiodic background is important for understanding neural spectra [5]. Sophisticated methods [5]–[7] support estimation of the 1/f-like, scale-free [8] or power-law [9] behavior of neural field spectra. Changes in *β*, the aperiodic exponent, have been investigated in many domains, including sleep [10]–[13], aging [14]–[16], and disease [17]–[19]. While many factors impact estimation of the aperiodic exponent (e.g., the frequency range analyzed [20], [21]), values of the exponent reported at higher frequencies (>20 Hz) typically range between −4 and −2 (Table 1).

**Table 1:** Example aperiodic exponents for human voltage spectra reported in the literature. The mean value of the aperiodic exponent (β) reported in the reference listed in column Ref. Additional details include the year of study, recording modality, number of subjects n and additional subject information, frequency range analyzed, and experimental condition. Considering only those studies with minimum frequencies >20 Hz (yellow highlighted rows), the aperiodic exponent has mean -3.1, lower quartile -4, and upper quartile -2.5.

| $\beta$ | Ref | Year | Recording modality | Frequency | Experimental condition |
| --- | --- | --- | --- | --- | --- |
| **-0.08** | [63] | 2019 | Scalp EEG (n=5) | 20-40 Hz | Anesthesia (Ketamine) |
| **-1.12** | [19] | 2022 | Scalp EEG (n=16, control subjects) | 1-40 Hz | Eyes closed |
| **-1.3** | [70] | 2021 | Scalp EEG (n=74, Tourette syndrome) | 2-40 Hz | Behavioral experiments |
| **-1.44** | [70] | 2021 | Scalp EEG (n=74, control) | 2-40 Hz | Behavioral experiments |
| **-1.48** | [19] | 2022 | Scalp EEG (n=18, stroke patients) | 1-40 Hz | Stroke patients |
| **-1.51** | [18] | 2019 | Scalp EEG (n=78, control subjects) | 4-50 Hz | Resting state |
| **-1.67** | [18] | 2019 | Scalp EEG (n=76, ADHD subjects) | 4-50 Hz | Resting state |
| **-1.84** | [10] | 2020 | Scalp EEG (n=9) | 30-45 Hz | Wakefulness |
| **-1.86** | [71] | 2013 | Scalp EEG (n=7, adults) | 0.2-30 Hz | Sleep |
| **-1.87** | [10] | 2020 | Scalp EEG (n=14) | 30-45 Hz | Resting state |
| **-2.03** | [63] | 2019 | Scalp EEG (n=5) | 20-40 Hz | Wakefulness |
| **-2.07** | [71] | 2013 | Scalp EEG (n=15, newborns) | 0.2-30 Hz | Sleep |
| **-2.32** | [72] | 2000 | Intracranial EEG (n=5) | 0.5-150 Hz | Resting state |
| **-2.33** | [12] | 2021 | Scalp EEG (n=175, T5) | 2-48 Hz | NREM sleep |
| **-2.44** | [31] | 2010 | Intracranial EEG (n=5) | 1-100 Hz | Wakefulness |
| **-2.48** | [63] | 2019 | Scalp EEG (n=5) | 20-40 Hz | Wakefulness |
| **-2.71** | [11] | 2022 | Scalp EEG (n=251) | 2-48 Hz | NREM sleep |
| **-2.73** | [12] | 2021 | Scalp EEG (n=175, Fz) | 2-48 Hz | NREM sleep |
| **-2.75** | [10] | 2020 | Intracranial EEG (n=12) | 30-45 Hz | Wakefulness |
| **-2.87** | [31] | 2010 | Intracranial EEG (n=5) | 1-100 Hz | Slow wave sleep |
| **-2.99** | [10] | 2020 | Intracranial EEG (n=10) | 30-45 Hz | Wakefulness |
| **-3.1** | [10] | 2020 | Scalp EEG (n=9) | 30-45 Hz | Anesthesia |
| **-3.13** | [63] | 2019 | Scalp EEG (n=5) | 20-40 Hz | Wakefulness |
| **-3.46** | [10] | 2020 | Scalp EEG (n=14) | 30-45 Hz | N3 Sleep |
| **-3.59** | [63] | 2019 | Scalp EEG (n=5) | 20-40 Hz | Anesthesia (Xenon) |
| **-3.67** | [10] | 2020 | Scalp EEG (n=14) | 30-45 Hz | N2 Sleep |
| **-3.69** | [10] | 2020 | Intracranial EEG (n=10) | 30-45 Hz | N3 Sleep |
| **-4** | [27] | 2009 | Intracranial EEG (n=20) | 80-500 Hz | Behavioral experiments |
| **-4.15** | [10] | 2020 | Intracranial EEG (n=10) | 30-45 Hz | REM sleep |
| **-4.34** | [10] | 2020 | Intracranial EEG (n=12) | 30-45 Hz | Anesthesia |
| **-4.36** | [63] | 2019 | Scalp EEG (n=5) | 20-40 Hz | Anesthesia (Propofol) |
| **-4.73** | [10] | 2020 | Scalp EEG (n=14) | 30-45 Hz | REM sleep |

The universal observation of 1/f-like neural field spectra with a restricted range of aperiodic exponents across different recording modalities, experiments, and neural systems suggests a common generative mechanism. However, many complex, specific mechanisms have been identified to generate this phenomenon. These include excitatory/inhibitory balance [22], low-pass frequency filtering by dendrites [23] or the extracellular medium [24], nonideal resistive components in the cell membrane [25], stochastic firing of neurons convolved with an exponential relaxation process [24], [26], [27], stochastic synaptic conductances [28], stochastically driven damped oscillators with different relaxation rates [29], local homogenous connectivity [30], combinations of many transient oscillations at different frequencies and amplitudes [31], or network mechanisms linking slower rhythms to broad neuronal recruitment and therefore larger amplitude field potentials [23]. Theoretically, scale-free phenomena – with 1/f-like behavior – have been linked to fractal properties [32], critical transitions [9], [33], and self-organized criticality [34], [35]. How these proposed biological and mathematical mechanisms contribute – separately or combined – to the range of aperiodic exponents observed across diverse neural field spectra remains unclear.

Here we demonstrate how, in general, a dynamical system with stochastic drive generates 1/f-like behavior at higher frequencies in neural field spectra. We show that two noise terms – representing correlated and uncorrelated noise inputs – produce the range of aperiodic exponents observed *in vivo*. We illustrate these general results in nonlinear models of neural and non-neural activity to demonstrate the ambiguity in determining the specific mechanisms given only the observed 1/f-like behavior in the spectrum. While more complex underlying mechanisms may exist, we propose instead that the range of aperiodic exponents observed *in vivo* is expected in general for dynamical systems with stochastic drive, without requiring a specific biological mechanism or tuning to collective critical behavior.

## RESULTS

As a general model of neural activity, we consider the n-dimensional dynamical system:

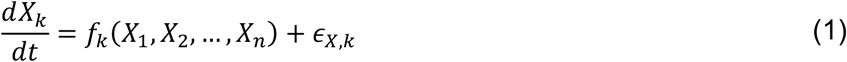

where, for each *k* = {1, 2, …, *n*}, *X*_*k*_ is a 1-dimensional variable, *f*_*k*_ is a nonlinear function, and *∈*_*x,k*_ is a noise term (Gaussian distributed with mean 0 and variance 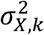). We assume the variable *X*_*1*_ (i.e., *k* = 1) is an observable quantity (e.g., the voltage recorded in the EEG or LFP) and all other variables (*X*_2_, *X*_3_, …, *X*_*n*_) represent n-1 unobserved or latent variables impacting the observable dynamics. The unspecified model (1) is general and therefore broadly consistent with diverse models of neural activity. To derive the main result from this general model requires no specific biophysical mechanism. In what follows, we illustrate these general results by making specific model choices for the variables (*X*_*k*_) and functions (*f*_*k*_).

We assume an equilibrium exists in the model so that 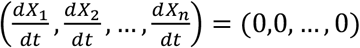 at 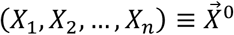. Near this equilibrium, the dynamics for one variable (*X*_*k*_) of the nonlinear system (1) can be approximated by the corresponding linear system,

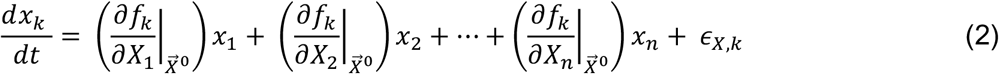

where *X*_*k*_ represent small deviations of the variable *k* from the equilibrium 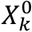, and we evaluate the partial derivatives of the nonlinear function *f*_*k*_ at the equilibrium 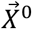 [36],[37]. To simplify notation, we express the system (2) as

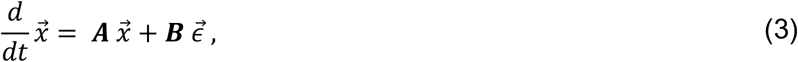

where 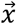 is the *n*-by-*n* vector of deviations from the equilibrium, ***A*** is the *n*-by-*n* Jacobian matrix of the nonlinear system (1) evaluated at the equilibrium, and ***B*** is an *n*-by-*n* diagonal matrix,

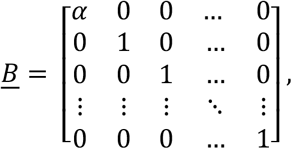

where the parameter *α* determines the noise level in the observable variable (*k* = 1) relative to the latent variables (*k* = {2, 3, …, *n*}), which we assume share the same noise level. The *n*-by-*n* vector 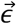 represents the ***A*** noise terms. We assume that ***A*** satisfies the conditions required for the linear system (3) to accurately approximate the dynamics of the nonlinear system (1) near the equilibrium (i.e., we assume ***A*** has no eigenvalues with zero real part and the equilibrium is therefore hyperbolic [36], [37]).

For the linear system (3), the cross-spectral matrix ***S***[*ω*] can be obtained from the expression [38]–[40],

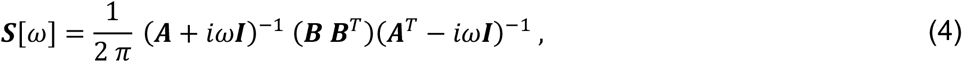

where ***A*** is real, ***I*** is the identity matrix, 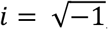, and *ω* = 2π*f* is the frequency. Evaluating the asymptotic behavior of the cross-spectral matrix at high frequencies (*ω* larger than any frequency associated with a natural rhythm of the linear system in (3)), the spectrum of the observable variable *X*_1_ is,

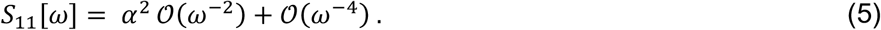

where 𝒪(*ω*^*k*^) indicates the limiting behavior of the spectrum as a function of the k^th^ power of *ω*; see Appendix. Without stochastic drive to the observable variable *X*_1_ (i.e., with *α* = 0),

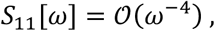

so that the aperiodic exponent *β* = −4 at high frequencies. Alternatively, with stochastic drive to the observable variable *X*_1_ (i.e., with *α* ≠ 0),

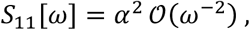

so that the aperiodic exponent *β* = −2 at high frequencies. We note that, in this case, random walk dynamics dominates the spectrum at high frequencies, with well-known power-law behavior (e.g., see [41]).

To summarize, we consider a general, n-dimensional nonlinear dynamical system with stochastic drive (1). We assume an equilibrium exists in this system with dynamics well-approximated by the linearized system (2). Near this equilibrium, the dynamics produce aperiodic exponents between −4 and −2 at high frequencies (i.e., frequencies beyond the natural frequencies or spectral peaks of the system) consistent with the values observed *in vivo*. The value of the aperiodic exponent depends on the relative noise in the observed and latent variables; when noise in the observable variable *X*_1_ dominates, the aperiodic exponent ≈ −2, while when the noise in the latent variables dominates, the aperiodic exponent ≈ −4.

The main result (5) and implications for the aperiodic exponent (−4 ≤ *β* ≤ −2) are for the general model (1). These general results do not require a specific biophysical model of neural activity. In what follows, we illustrate the generality of these results in four example models, in which we choose the nonlinear functions *f*_*k*_ in model (1). In doing so, we show that each model produces aperiodic exponents consistent with *in vivo* data (−4 ≤ *β* ≤ −2) but with different physical interpretations.

### Example 1: The Wilson-Cowan model

We first consider the Wilson-Cowan equations as a model of neural population activity [42]. The equations describe the interacting dynamics of an excitatory (*E*) and inhibitory (*I*) neural population,

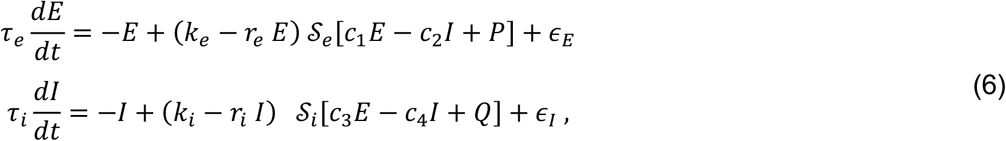

where *𝒮* is the sigmoid function,

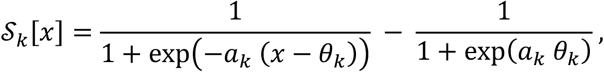

and *∈*_*E*_ and *∈*_*I*_ are noise terms (Gaussian distributed with mean 0 and variances 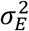 and 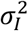, respectively). We choose the parameters to produce damped oscillatory behavior (see Figure 10 of [42] and the caption of Figure 1 for the parameter values used here). With these parameters, we find an equilibrium of the model (6),

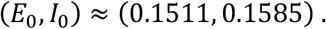

The linearized system near this equilibrium is approximately,

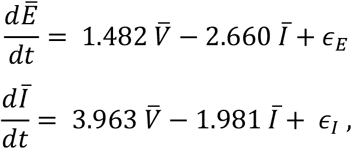

and the equilibrium is hyperbolic, with eigenvalues −0.250 ± 2.75 *i*.

**Figure 1:**
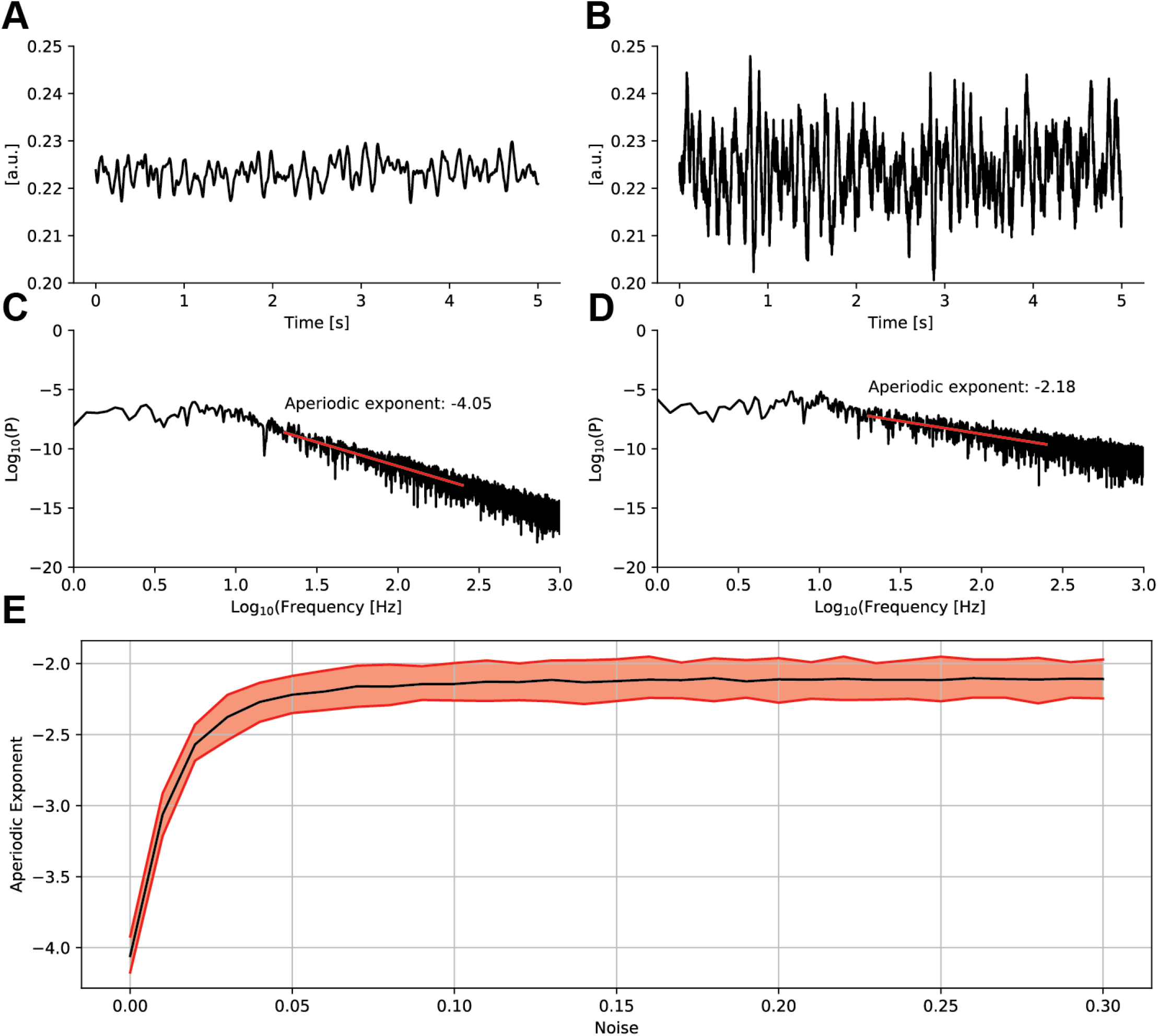
In the Wilson-Cowan model, the aperiodic exponent increases from approximately −4 to −2 with the excitatory noise. **(A**,**B)** Example excitatory population time series when (A) the excitatory noise is 0 (i.e., ∈_E_ = 0), or (B) the excitatory noise is non-zero (i.e., α_E_ = 0.2). **(C**,**D)** The corresponding spectra (black) and linear fits (red, 20 Hz to 250 Hz) for the time series in (A,B). **(E)** Estimates of the aperiodic exponent for increasing values of excitatory noise. Black (red) indicates mean (standard deviation) of estimates across 100 simulations. In all simulations, the inhibitory noise (α_1_ = 0.1) is fixed. We use the model parameters: c_1_ = 15, c_2_ = 15, c_3_ = 15, c_4_ = 7, a_e_ = 1, λ_e_ = 2, a_i_ = 2, λ_i_ = 2.5, τ_e_ = 50, τ_i_ = 50, r_e_ = 1, r_i_ = 1, k_e_ = 1, k_i_ = 1, P = 1.25, Q = 0. Code to simulate the model and create this figure is available at https://github.com/Mark-Kramer/Aperiodic-Exponent-Model.

According to the general theory, we expect the nonlinear system (6) near the equilibrium (*E*_0_,*I*_0_) to produce spectra with aperiodic exponents −4 ≤ *β* ≤ −2, depending on the values of the stochastic drives (*∈*_*E*_, *∈*_*I*_). To show this, we simulate the nonlinear system (6) and estimate the spectrum of the excitatory variable *E* with fixed inhibitory noise (σ_*I*_ = 0.1) and variable excitatory noise (0 ≤ σ_*E*_ ≤ 0.3). In agreement with the general theory (Figure 1), as the excitatory noise increases, the aperiodic exponent increases from near *β* ≈ −4 when σ_3_ = 0 to *β* ≈ −2 when σ_3_ = 0.3.

We conclude that this nonlinear model of neural population activity (6) produces aperiodic exponents consistent with the range of values observed *in vivo*. In this model, the value of the aperiodic exponent depends on the relative noise in the observable variable *E* and latent variable *I*. When noise in the inhibitory population dominates, the aperiodic exponent approaches −4; when noise in the excitatory population dominates, the aperiodic exponent approaches −2.

### Example 2: A Hodgkin-Huxley type model

We next consider a reduced Hodgkin-Huxley type model [36], [43]. We choose this model to simulate the subthreshold dynamics of a single neuron, in contrast to the neural population dynamics simulated in the previous example. The model equations describe the dynamics of an observable voltage (*V*) and a latent membrane current (*M*),

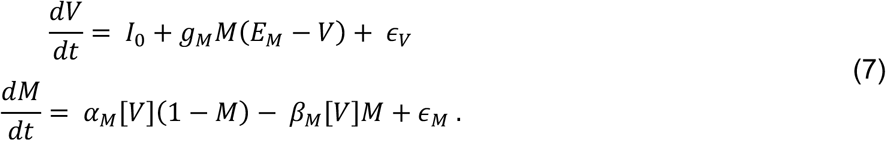

In the first equation, three terms drive the voltage dynamics: a constant input current (***I***_0_), a dynamic membrane current (*g*_*M*_*M*(*E*_*M*_ − *V*)), and voltage noise (*σ*_*V*_, Gaussian distributed with mean 0 and variance 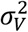). In the second equation, the dynamics of the latent membrane current depend on forward (*α*_*M*_[*V*]) and backward (*β*_*M*_[*V*]) rate functions,

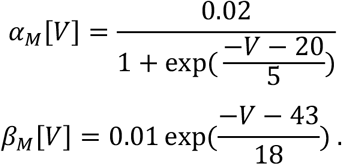

We choose these functions to simulate a muscarinic receptor suppressed potassium current (M-current, see Table A2 of [44]) [45], [46]. We omit other membrane currents (e.g., fast sodium and potassium currents) to focus on the subthreshold membrane dynamics without action potential generation. A stochastic drive also impacts the membrane current dynamics (*∈*_*M*_, Gaussian distributed with mean 0 and variance 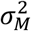).

Choosing the model parameters *I*_0_ = 1, *g*_*M*_ = 4, and *E*_*M*_ = −95, we find an equilibrium of the model (7) at

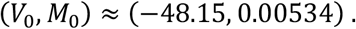

The linearized system near this equilibrium is approximately,

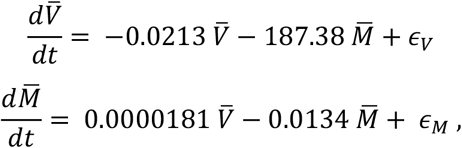

and the equilibrium is hyperbolic with eigenvalues −0.0174 ± 0.0581 *i*.

Consistent with the general theory, we expect this nonlinear Hodgkin-Huxley type model (7) to produce spectra with aperiodic exponents −4 ≤ *β* ≤ −2, depending on the values of the stochastic drives (*∈*_*V*_, *∈*_*M*_). To show this, we simulate the Hodgkin-Huxley type model (7) and estimate the spectrum of the voltage variable *V* with fixed current noise (*σ*_*M*_ = 0.01) and variable voltage noise (0 ≤ σ_*V*_ ≤ 30). In agreement with the general theory (Figure 2), as the voltage noise increases, the aperiodic exponent increases from near *β* ≈ −4 when σ_*V*_ = 0 to *β* ≈ −2 when σ_*V*_ = 30.

**Figure 2:**
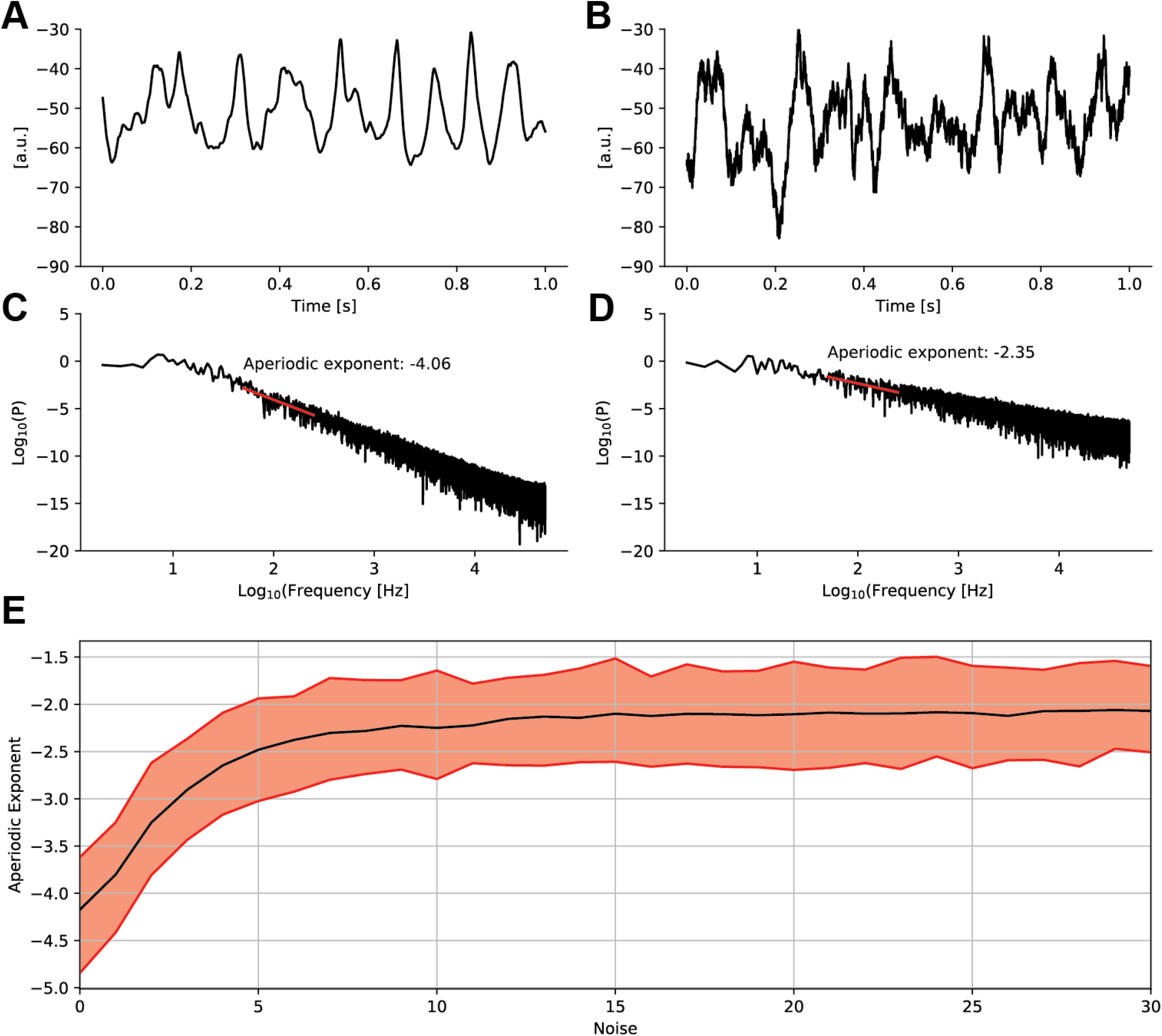
In a Hodgkin-Huxley type model, the aperiodic exponent increases from approximately −4 to −2 with the voltage noise. **(A**,**B)** Example voltage time series when (A) the voltage noise is 0 (i.e., ∈_V_ = 0), or (B) the voltage noise is non-zero (i.e., σ_V_ = 20). **(C**,**D)** The corresponding spectra (black) and linear fits (red, 50 Hz to 250 Hz) for the time series in (A,B). **(E)** Estimates of the aperiodic exponent for increasing values of voltage noise. Black (red) indicates mean (standard deviation) of estimates across 100 simulations. In all simulations, the current noise (σ_M_ = 0.01) is fixed. Code to simulate the model and create this figure is available at https://github.com/Mark-Kramer/Aperiodic-Exponent-Model

We conclude that this nonlinear model of single neuron subthreshold dynamics (7) produces aperiodic exponents consistent with the range of values observed *in vivo*. In agreement with the general theory, the value of the aperiodic exponent depends on the relative noise in the observable voltage variable and latent current variable. In this case, when noise in the membrane current dominates, the aperiodic exponent approaches −4; when noise in the voltage dominates, the aperiodic exponent approaches −2.

### Example 3: A model of predator-prey interactions

To illustrate the generality of the main result (5) beyond models of neural activity we consider a third nonlinear model of predator-prey interactions,

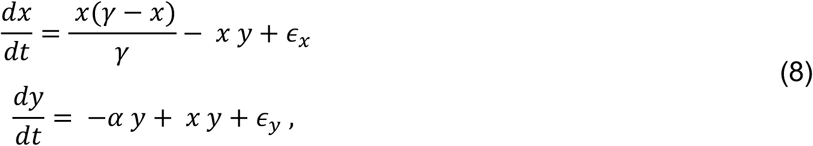

where *x* and *y* represent prey and predator populations, respectively, and the prey population includes self-regulation; (*α, γ*) are positive constants; and *∈*_*x*_ and *∈*_*y*_ are noise terms (Gaussian distributed with mean 0 and variances 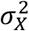 and 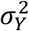, respectively) [47]. The non-trivial equilibrium of the deterministic system is,

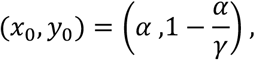

and we require *γ* > *α* so that the equilibrium predator population is positive. The linearized system near this equilibrium is,

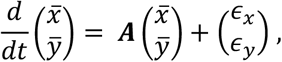

where 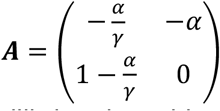. Because the trace of 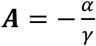 is negative, and the determinant of ***A*** = *αy*_0_ is positive, the equilibrium is stable and hyperbolic [47].

Consistent with the general theory, we expect this predator-prey model (8) to produce spectra with aperiodic exponents −4 ≤ *β* ≤ −2, depending on the values of the stochastic drives (*∈*_*x*_, *∈*_*y*_). To show this, we simulate the predator-prey model (8) over a range of parameters (*α, γ*) and estimate the spectrum of the prey variable *x* with fixed predator noise (σ_*y*_ = 1) and variable prey noise (0 ≤ σ_*x*_ ≤ 0.005). In agreement with the general theory, the values of the aperiodic exponent in this non-neural model lie within the range −4 ≤ *β* ≤ −2, depending on the relative noise in the prey variable (Figure 3). As the prey noise increases, the aperiodic exponent increases from near *β* ≈ −4 when σ_*x*_ = 0 to *β* ≈ −2 when σ_*x*_ = 0.005 across a range of model parameters (*α, γ*).

**Figure 3:**
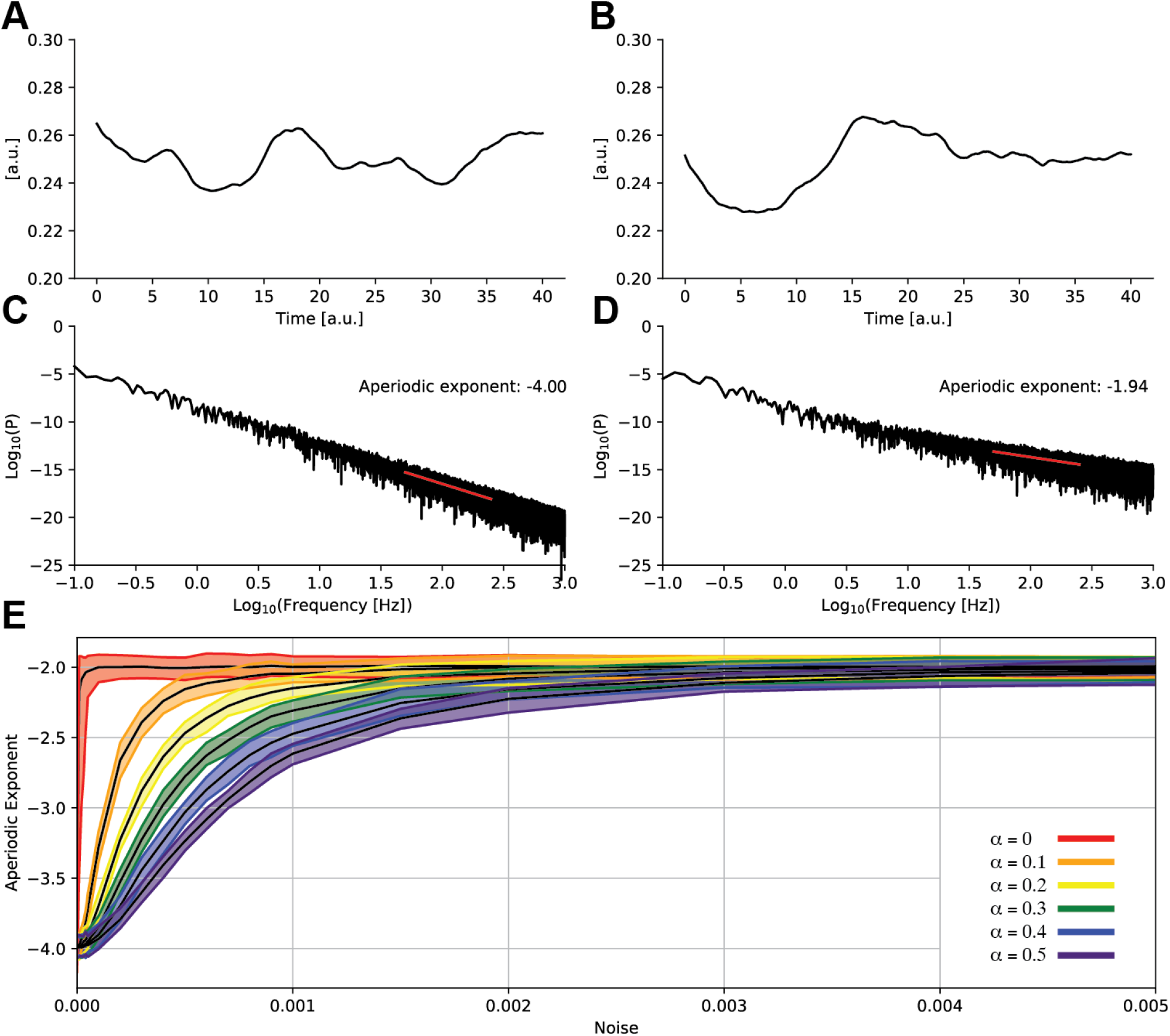
In a predator-prey population model, the aperiodic exponent increases from approximately −4 to −2 with the prey noise. **(A**,**B)** Example prey population dynamics when (A) the prey noise is 0 (i.e., α_x_ = 0), or (B) the prey noise is non-zero (i.e., σ_x_ = 0.01). **(C**,**D)** The corresponding spectra (black) and linear fits (red, 50 Hz to 250 Hz) for the time series in (A,B). **(E)** Estimates of the aperiodic exponent for increasing values of prey noise. Black (color) indicates mean (standard deviation) of estimates across 100 simulations. In all simulations, the predator noise (σ_y_ = 1) is fixed and the model simulated for 250,000 steps with sampling interval (dt = 0.002; to avoid an initial large amplitude transient, we omit the first 50,000 steps of simulated data from analysis. In (A-D), the parameters are α = 0.25, γ = 0.6. In (E), colors indicate simulations at α = {0.0, 0.1, …, 0.5}. For each fixed α, we simulate 10 instances of the model at 10 different values of γ = {0.1, 0.2, 0.3, …, 1.0} for a total of 100 simulations. Code to simulate the model and create this figure is available at https://github.com/Mark-Kramer/Aperiodic-Exponent-Model

### Example 4: A 10^th^-order model of macroscopic voltage fluctuations

Finally, to illustrate an application of the main result (5) to a higher-dimensional model, we consider a mean-field model of neural population activity consisting of the coupled stochastic differential equations:

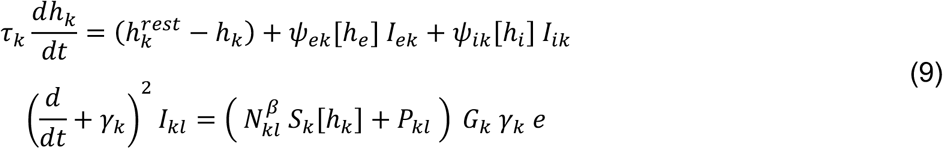

where *k* = {*e, i*} and *l* = {*e, i*} denote excitatory (*e*) and inhibitory (*i*) neural populations, *ψ*_*kl*_ are normalized weighting functions, and ***S***_*k*_[*h*_*k*_] are sigmoidal transfer functions [48]. The model variables simulate the macrocolumn-averaged transmembrane soma voltage of an excitatory (*h*_*e*_) and inhibitory (*h*_*i*_) neural population, and synaptic input (***I***_*kl*_) from population *k* to population *l*. Expressing the second-order differential equations for the synaptic inputs ***I***_*kl*_ as first-order differential equations results in a system of 10 coupled first-order differential equations. The variable *h*_*e*_ is observable [49] and the other variables are latent. We include stochastic drive to the observable variable (mean 0 and variance 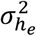) and each latent variable (mean 0 and variance 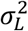 to all latent variables). Applications of the model include simulating electroencephalogram (EEG) dynamics during sleep [50]–[52], seizures [53]–[56], and anesthesia [48], [49].

Fixing all model parameters to the default values in [48], a stable equilibrium exists [48]. We therefore expect, consistent with the general theory, the model (9) to produce spectra with aperiodic exponents −4 ≤ *β* ≤ −2, depending on the values of the stochastic drives. To show this, we simulate the model (9) and estimate the spectrum of the voltage variable *h*_*e*_ with fixed noise (*α*_*L*_ = 50) to all latent variables and variable noise to the observable variable 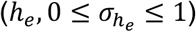. In agreement with the general theory (Figure 4), as the noise to the excitatory neural population increases, the aperiodic exponent increases from near *β* ≈ −4 when 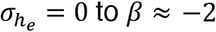 when 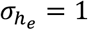.

**Figure 4:**
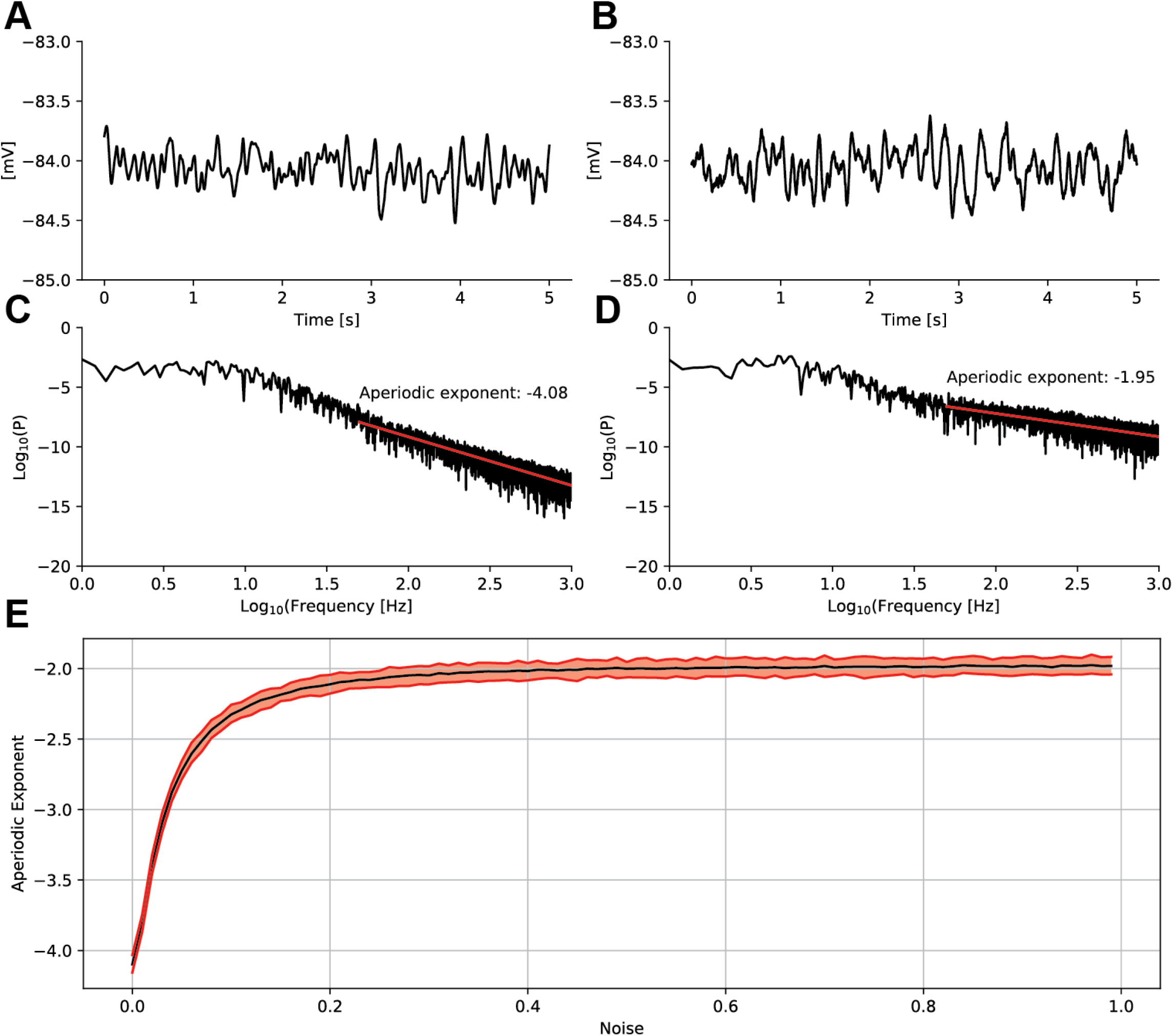
In a 10^th^ order mean-field model of neural population activity, the aperiodic exponent increases from approximately −4 to −2 with increasing noise to the excitatory neural population. **(A**,**B)** Example excitatory neural population activity when (A) noise to the excitatory neural population is 0 (i.e., 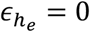), or (B) non-zero (i.e., 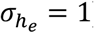). **(C**,**D)** The corresponding spectra (black) and linear fits (red, 50 Hz to 1000 Hz) for the time series in (A,B). **(E)** Estimates of the aperiodic exponent for increasing values of excitatory neural population noise. Black (red) indicates mean (standard deviation) of estimates across 100 simulations. In all simulations, the latent noise terms (σ_L_ = 50) are fixed. Code to simulate the model and create this figure is available at https://github.com/Mark-Kramer/Aperiodic-Exponent-Model

We conclude that this high-dimensional (10^th^ order) nonlinear model of macroscopic neural population activity (9) produces aperiodic exponents consistent with the range of values observed *in vivo*. In agreement with the general theory, the value of the aperiodic exponent depends on the relative noise in the observable variable (*h*_*e*_) and latent variables. In this case, when noise outside of the excitatory population dominates, the aperiodic exponent approaches −4; when noise in the excitatory population (*h*_*e*_) dominates, the aperiodic exponent approaches −2.

The four example simulations demonstrate the difficulty of interpreting the biological mechanism producing the aperiodic exponent. While the four models (6-9) simulate different neural and non-neural dynamics, each model produces 1/f-like scaling with aperiodic exponents consistent with *in vivo* observations of neural activity. The interpretation of the mechanism affecting the aperiodic exponent depends on the model choice; the aperiodic exponent *β* approaches −2 with increasing noise to an excitatory neural population in (6) and (9), voltage noise in (7), or prey noise in (8). Because each model satisfies the general conditions (i.e., a hyperbolic equilibrium exists), the spectral results derived for the general model in (5) capture the 1/f-like spectrum for each specific, biophysical model considered here.

## DISCUSSION

A universal feature of neural field potential spectra is 1/f-like scaling. To produce this power-law, many generative mechanisms have been proposed with diverse biological implementations and interpretations. Here, we do not propose a specific biological mechanism produces 1/f-like scaling. Instead, to understand how different mechanisms can produce similar 1/f-like scaling, we consider a general nonlinear dynamical system with stochastic drive. We show that dynamics near a (hyperbolic) equilibrium in this general model produce aperiodic exponents between −4 and −2, consistent with the range of values reported *in vivo* for higher frequencies (e.g., >20 Hz). We illustrate these results in neural and non-neural models. We conclude that the range of aperiodic exponents observed across recording modalities, experiments, and neural systems is a natural consequence of a noise driven dynamical system.

We considered here a single statistic – the aperiodic exponent – reflecting the 1/f-like feature of the neural field spectrum. We note that many different models can explain the same observed statistic. For example, observations from neural systems can produce spectra with broadband peaks in the gamma band (approximately 30-80 Hz) [57]. Many approaches exist to explain these observed spectral peaks, including statistical approaches (e.g., an autoregressive model of order two [58]), mechanical approaches (e.g., a damped driven oscillator [58]), or biophysical approaches (e.g., the interneuron network model, or the pyramidal-interneuron network model [59]). Our understanding of the gamma rhythm in a particular experiment depends on the model choice. In the same way, observations from neural systems produce spectra with 1/f-like scaling, and many models exists to explain this scaling. Our results show mathematically why many models can produce the range of aperiodic exponents observed *in vivo*(−4 ≤ *β* ≤ −2). Due to the general nature of this mathematical result, it is not surprising that many different proposals exist to explain the 1/f-like spectrum. In practice, we expect biological mechanisms must exist to create the scaling observed *in vivo*. Identifying these biological mechanisms, and their expression in the diverse observations of 1/f-like scaling reported, remains an important challenge.

Our results are consistent with previous work showing that power-law scaling occurs in simple stochastic or physical systems. For example, in [22] the authors relate estimates of the aperiodic exponent to changes in the balance between excitation and inhibition. To do so, the authors simulate the local field potential as the summed synaptic current generated by independent stochastic spiking excitatory and inhibitory cells. In [24] the authors propose that 1/f-like scaling in the spectrum does not rely on critical states, but instead depends on the filtering properties of the extracellular medium (although this mechanism remains debated [60]). In [61] the authors show that simple models of stochastic processes (high-frequency shot-noise processes or Ornstein-Uhlenbeck processes) produce peak-amplitude distributions consistent with power-law distributions. In [62] the authors show that inhomogeneous Poisson processes can produce approximate power law distributions in the size and duration distributions of avalanches (i.e., activity cascades). Consistent with our results, these examples generate power-low scaling without requiring a sophisticated biological or mathematical mechanism. Distinct from these previous works, we consider an (unspecified) n-dimensional dynamical system and show that, near a (hyperbolic) equilibrium, stochastic drive produces aperiodic exponents consistent with values observed *in vivo*.

Under the general framework considered here, the range of aperiodic exponents reflects different types of noise. We note that the observable dynamics (e.g., variable *X*_1_ in (1)) depend on multiple types of noise: directly on the stochastic drive to the observable variable, and indirectly on the stochastic drive to the latent variable(s). The latent dynamics introduce correlations in the uncorrelated latent noise process before this noise reaches the observable dynamics. Therefore, the observable dynamics depend on both uncorrelated and correlated noise inputs, and we may interpret the aperiodic exponent in terms of these different types of noise; an aperiodic exponent near *β* = −4 indicates correlated noise inputs dominate the high frequency observable dynamics, while *β* = −2 indicates uncorrelated noise dominates the high frequency observable dynamics. These general results are independent of a specific biophysical mechanism, and a biophysical interpretation of the relationship between different types of noise and the aperiodic exponent depends on the specific model choice.

Some observations report aperiodic exponents greater than −2 (i.e., β > −2), beyond the range of aperiodic exponents derived here. Experimental factors, such as measurement noise, which flattens the spectrum and shifts β towards 0, might contribute to these observations. In addition, analysis factors may impact reported results. For example, the frequency range considered varies widely across studies (see discussion of fitting ranges in [21]). In general, larger aperiodic exponents (more negative *β*) occur at higher frequencies [24], [63]– [65], although not always [66]. In the framework considered here, the result −4 ≤ *β* ≤ −2 holds in the high frequency limit (when the measured frequency exceeds any natural frequency of the neural population) and near an equilibrium of the dynamical system. We expect analyzing lower frequency bands (e.g., below the natural frequency), or analyzing the low and high frequency bands after accounting for the broadband peak, will increase the aperiodic exponent. While potentially useful, the aperiodic exponent reported in lower frequency bands is more difficult to interpret. To assess low frequency rhythms requires long durations of data, which increases the chance of nonstationarity. Artifacts (e.g., slow drifts) and analysis choices (e.g., whether to subtract the signal mean) also impact the low frequency power. Finally, while power-law features at low frequencies may reflect the same power-law features at high frequencies, these different phenomena unlikely reflect the same neural mechanisms. We also expect analyzing dynamics away from an equilibrium of the dynamical system will increase the aperiodic exponent. Away from an equilibrium, nonlinear terms in the model have a greater impact on the dynamics and resulting spectrum. These nonlinearities may increase power at high frequencies (e.g.,[67]) and therefore increase the aperiodic exponent beyond the range derived for the linear dynamics near the equilibrium.

Many well-supported observations of power-laws appear in neuroscience (e.g., avalanches of population voltage discharges [68], amplitudes of narrowband oscillations [69]). Here, we consider one type of power-law: the 1/f-like neural field spectrum and a general, noise-driven dynamical system. Under this general model, the aperiodic exponent represents the impact of noise in the observable and latent dynamics, without requiring a sophisticated biological or dynamical mechanism. We propose that the range of aperiodic exponents −4 ≤ *β* ≤ −2 observed *in vivo* represents the expected dynamics near an equilibrium in a nonlinear dynamical system driven by noise. The generality of the model is consistent with the universality of 1/f-like field spectra, reflecting a basic dynamical feature present in many different neural systems. However, this simplicity may also limit the computational utility of this mechanism and the role of the aperiodic exponent in measuring neural computations.

## MATERIALS AND METHODS

### Estimation of the aperiodic exponent β from time series data

To estimate the aperiodic exponent *β*, we first compute the spectrum in the standard way. To a simulated voltage time series (*V*_*t*_) with sampling interval Δ and duration *T*, we subtract the mean, apply a Hanning taper, compute the Fourier transform (*V*_*f*_), and multiply by the complex conjugate 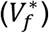:

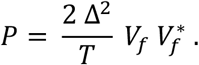

We note that the square of the Fourier coefficients is essential for consistent interpretation of the aperiodic exponent across studies; omitting the square is a common mistake identified in previous work (see discussion in [41]). For frequencies *f*, we fit a linear model to the logarithm base 10 of the spectrum (log_10_ *P*) with predictor logarithm base 10 of the frequency (log_10_ *f*),

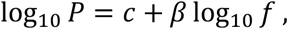

where *β* is the estimate of the aperiodic exponent. Code to compute the spectrum and estimate the aperiodic exponent is available at https://github.com/Mark-Kramer/Aperiodic-Exponent-Model.

## ACKNOWLEDGEMENTS

The authors would like to thank Professor Konstantinos Spiliopoulos and Professor Uri Eden for recommendations regarding the analysis of stochastic differential equations. The authors were supported in part by NSF #1451384, NIH R01NS110669, and NIH R01NS119483.

## APPENDIX

### Derivation of the asymptotic behavior of the cross-spectral matrix

Consider the two terms of (4) that involve the Jacobian,

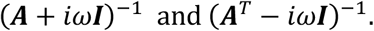

In general, these *n*-by-*n* matrices are complicated expressions of the constants in ***A*** and powers of *ω*. To characterize the limiting behavior of these matrices for large values of *ω*, we express each of these two terms using asymptotic notation,

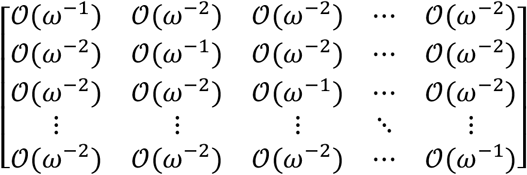

where terms on the diagonal grow proportional to *ω*^-1^ and terms off the diagonal grow proportional to *ω*^-2^. The expression

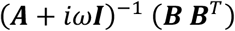

in (4) then becomes

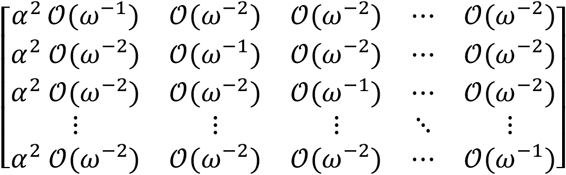

and the cross-spectral matrix ***S***[*ω*] in (4)

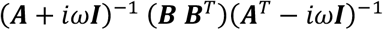

becomes

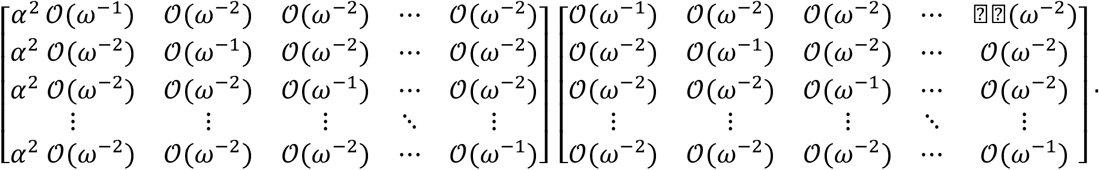

The spectrum of the observable variable (***S***_11_[*ω*]) corresponds to the first entry of this matrix product,

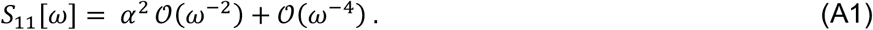

To illustrate these general results, we consider as a specific example the 2-dimensional dynamical system with\

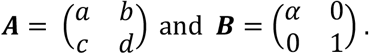

Evaluating the cross-spectral matrix (4) for the observable variable (***S***_*XX*_[*ω*]) we find,

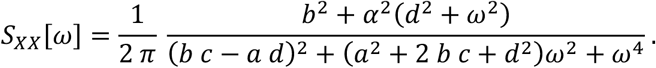

Isolating the *α*^2^ term in the numerator, this expression becomes,

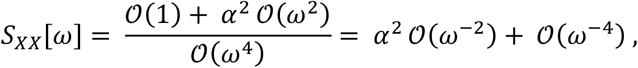

equivalent to the general expression (A1).

